# Global Cancer Transcriptome Quantifies Repeat Element Polarization Between Immunotherapy Responsive and T cell Suppressive Classes

**DOI:** 10.1101/145946

**Authors:** Alexander Solovyov, Nicolas Vabret, Kshitij S. Arora, Alexandra Snyder, Samuel A. Funt, Dean F. Bajorin, Jonathan E. Rosenberg, Nina Bhardwaj, David T. Ting, Benjamin D. Greenbaum

**Affiliations:** Tisch Cancer Institute, Departments of Medicine, Hematology, and Medical Oncology, Icahn School of Medicine at Mount Sinai, New York, New York, USA.; Department of Oncological Sciences, and Department of Pathology, Icahn School of Medicine at Mount Sinai, New York, New York, USA.; Joint first author; Massachusetts General Hospital Cancer Center; Department of Pathology, and; Department of Surgery, Harvard Medical School, Charlestown, Massachusetts, USA; Department of Medicine, Memorial Sloan Kettering Cancer Center, New York, New York, United States of America; Department of Medicine, Weill Cornell Medical College, New York, New York, United States of America; Department of Medicine, Harvard Medical School, Charlestown, Massachusetts, USA; Joint corresponding author

## Abstract

Growing evidence indicates that innate immune pathway activation is critical for responses to immunotherapy and overall cancer prognosis. It has been posited that innate immunity in the tumor microenvironment can be driven by derepression of endogenous repetitive element RNA. The ability to characterize these species can potentially provide novel predictive biomarkers for tumor immune responses and a mechanistic basis for elements of innate activation by tumors. We first compared total RNA and poly(A)-capture protocols applied to tumor RNA-sequencing to detect non-RNA coding transcriptomes in the TCGA. While the poly(A) protocol efficiently detects coding, most non-coding genes, and much of the LINE/SINE/ERV repeat repertoire, we found that it fails to capture overall repeat expression and co-expression. The probing of total RNA expression reveals distinct repetitive co-expression subgroups. Secondly, we found that total repeat element expression delivers the most dynamic changes in samples, which may serve as more robust biomarkers of clinical outcomes. Finally, we show that while expression of ERVs, but not other immunostimulatory repeats such as HSATII, is associated with response to immunotherapy in a cohort of patients with urothelial cancer treated with anti-PD-L1 therapy, global repeat derepression strongly correlates with an immunosuppressive phenotype in the microenvironment of colorectal and pancreatic tumors. We validate *in situ* in human primary tumors, associating the immunosuppressive phenotype with HSATII expression. In conclusion, we demonstrate the importance of analyzing repetitive element RNAs as potential biomarkers of response to immunotherapy and the need to better characterize these features in next generation sequencing protocols.

## Introduction

The transcriptional landscape of a cancer cell extends well beyond protein-coding messenger RNA and includes numerous non-coding transcripts, some of which play essential roles in modulating malignant transformation (Lin and He 2017). Among the different classes of noncoding RNA (ncRNA) are repetitive elements, which comprise more than half of the human genome and were shown to undergo increased transcriptional activity during neoplasia (Ting et al. 2011; Criscione et al. 2014). Aberrant transcription of repetitive elements in tumors is likely modulated by epigenetic modifications (Carone and Lawrence 2013) and loss of tumor suppressor function (Wylie et al. 2015, Levine et al. 2016). Moreover, many repeat RNAs include sequence and structure patterns typically found in pathogen genomes rather than human genomes (Chiappinelli et al. 2015, Roulois et al. 2015, Tanne et al. 2015). Such pathogen “mimicry” can be detected by innate pattern recognition receptors (PRRs), initiating signaling in the tumor microenvironment relevant for immune and epigenetic therapies (Leonova et al. 2013, Chiappinelli, et al. 2015, Roulois et al. 2015, Woo et al. 2015; Desai et al. 2017, Greenbaum 2017). These direct immunomodulatory features of repetitive elements provide a novel functional signaling pathway not previously appreciated in human cancers.

Unfortunately, a practical barrier to assessing the landscape of aberrantly transcribed immunostimulatory repetitive elements has been the typical protocols employed in next generation RNA sequencing (RNA-seq) (Zhao et al. 2014). The vast majority of publicly available RNA-seq datasets contain only sequences of polyadenylated RNA and, as we show, such approaches fail to detect many putatively functional non-coding transcripts that can stimulate PRRs. To give a sense of the degree to which protocols are biased in this regard, one needs only to look at the statistics of The Cancer Genome Atlas (TCGA). While thousands of solid tumors are sequenced using the poly(A) select approach, only 38 solid tumor samples probe the total RNA. The breadth of aberrant repetitive element transcription and its link to pathogen mimicry in cancer is therefore severely under-quantified despite their great potential importance as cancer biomarkers and causal agents of innate immune activation.

In this work, we first examined the 29 samples from TCGA where both poly(A) enriched and total RNA-seq data are available from the same tumor. We find a large amount of missing repetitive element transcripts from tumors sequenced using poly(A) protocols. Second, we show that repetitive elements expressed from tumors fall into a set of distinct co-expression clusters. These clusters correspond to common repeat classes, such as long interspersed nuclear elements (LINEs), satellite repeats, and endogenous retroviruses (ERVs). We quantify the nature of these clusters, their diversity, and whether they have anomalous motif usage, indicating a potential to trigger PRRs. We unravel new associations between expression of specific classes of repetitive elements, patient survival rates and the immune profile of the tumor microenvironment. In doing so, we find a relationship between ERV expression in patients with urothelial cancer treated with PD-L1 blockade and response to the therapy, and an immunosuppressive phenotype associated with global depression in pancreatic and colon cancers.

## Results

### Proper normalization of total and poly(A) selected sequencing shows widespread differences in repetitive element detection

We identified 29 patient samples in TCGA that had RNA-seq data prepared using both the poly(A) enriched and total RNA protocol. Gene expression values computed from total RNA and poly(A) sequencing cannot be compared directly due to gene specific biases inherent to each protocol. These gene specific biases cannot be corrected using standard normalization methods, unlike sample specific bias. However, we find that by applying trimmed mean of M-values (TMM) normalization (Robinson and Oshlack 2010) between 29 paired patient samples - and clustering samples based on protein coding genes only - the same patient’s samples will cluster together, despite having different sequencing library construction protocols (Fig. 1A, black/white color code at the top). The technical difference between the poly(A) and the total RNA protocol is therefore less than the biological difference for protein coding genes in the 29 patients in our cohort. A similar picture (but to a lesser extent) was observed when we examined the computed expression of annotated non-coding RNAs (Fig. 1B). Evaluation of repetitive element expression, however, was markedly different between total and poly(A) RNA protocols. For most repeats, expression computed using the total RNA protocol exceeded that computed from the poly(A) protocol (Fig. 1C). This is true even if one compares expression from unrelated patients. Hierarchical clustering for repeat expression is therefore completely governed by the protocol used to prepare the RNA-seq library.

**Figure 1.**
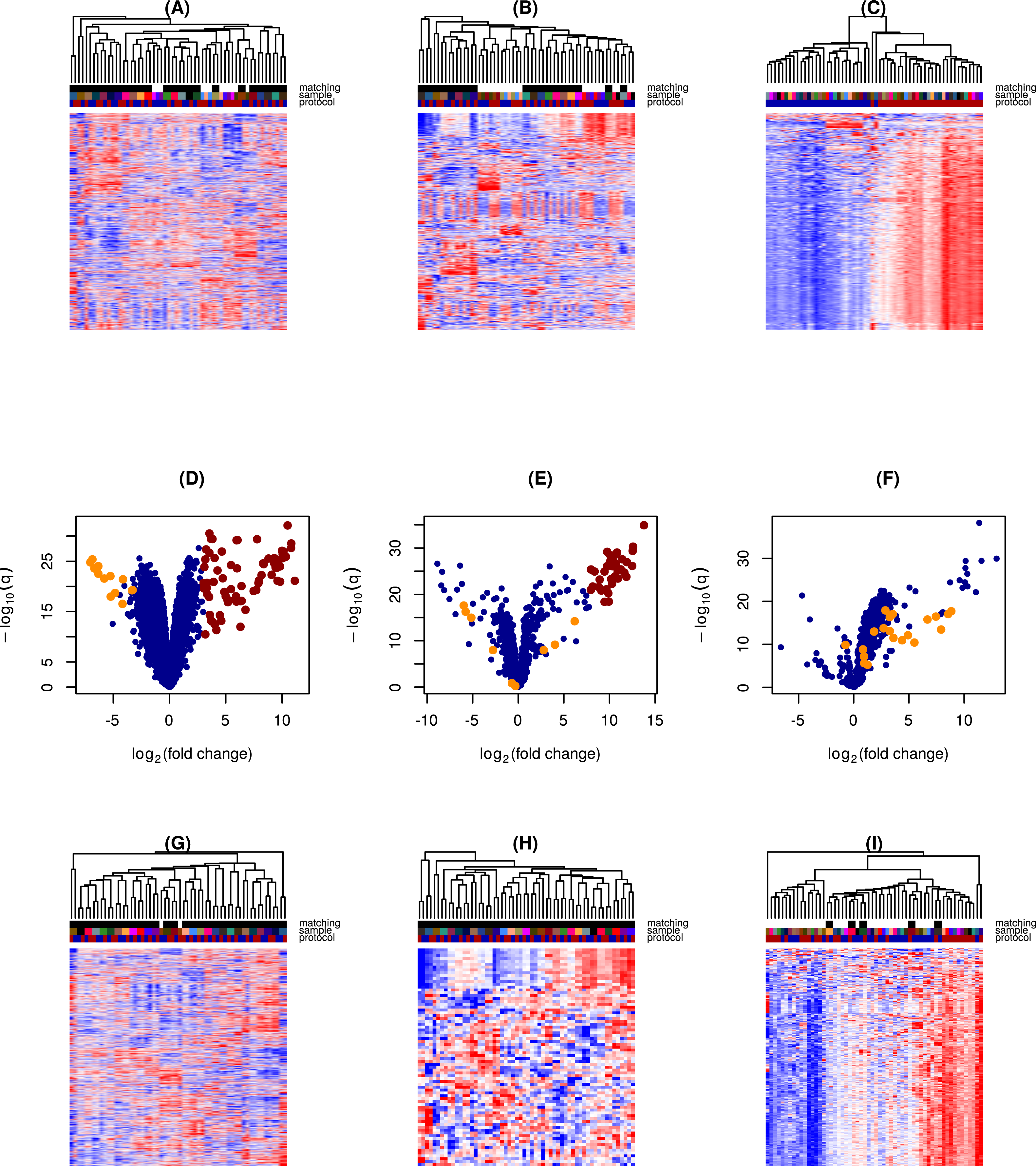
(A), (B), (C) Hierarchical clustering and expression heat map based on the coding gene expression (A), non-coding gene expression (B) and repeat element expression (C). Color code at the top: total RNA and poly(A) prepared aliquots from the same sample are denoted with the same color. Total RNA protocol is denoted by the red color, poly(A) protocol is denoted by the blue color. The black/white color at the top indicates whether the total RNA and the poly(A) aliquots were direct neighbors in the dendrogram. The total RNA and the poly(A) enriched aliquots were the direct neighbors in the dendrogram for 23 out of 29 pairs for coding genes and 18 out of 29 pairs for non-coding genes. The clustering for the repeat elements is governed by the preparation protocol. There are the two subclusters corresponding to the poly(A) and the total RNA aliquots. , (E), (F) Volcano plots for the pairwise difference in the computed expression between the poly(A) and the total RNA protocol. Positive log(fold change) means the higher computed expression in the total RNA protocol. Both coding (D) and non-coding (E) genes exhibit different biases (i.e., positive of negative log (fold change)) with a few outliers. Mitochondrial genes (shown in orange) are depleted in the total RNA protocol. Computed expression of repeat elements (F) is higher in the total RNA protocol for all but a few elements. See also Table S1. (G), (H), (I) Hierarchical clustering and expression heat map based on the adjusted coding gene expression (G), non-coding gene expression (H) and repeat element expression (I). Only genes detectable (i.e., having a sufficient read number) in both protocols are included. In the absence of the technical noise the computed expression difference between the two protocols would be a gene specific sample independent constant.

Out of 13740 coding genes, 2615 (19%) had significantly lower computed expression (FDR < 0.05 and fold change > 1.5) and 2256 (16%) had significantly higher computed expression in the total RNA protocol. Out of 893 annotated non-coding genes, 228 (25%) had significantly lower computed expression and 197 (22%) had significantly higher computed expression in the total RNA protocol. Out of 967 repeat elements, 31 (3%) had significantly lower computed expression and 831 (86%) had significantly higher computed expression in the total RNA protocol. Interestingly, some coding genes (75 out of 13740, 0.5%) form an outlier population with higher computed expression in the total RNA protocol (Fig. 1D). Those were mostly histone related genes on chromosome 6. For non-coding genes, 38 out of 893 (4%) were such outliers mostly composed of small RNAs (17 snoRNA, 7 misc_RNA, 6 snRNA, 5 scaRNA, 2 antisense, 1 miRNA) (Fig. 1E). Conversely, in the case of repeats, there is a clear and consistent inability to capture repetitive element expression using the poly(A) protocol (Fig. 1F).

If the effect of preparation on computed gene expression is sample independent, expression computed from paired total RNA and poly(A) samples will differ by a gene specific constant independent of the sample. We designed an analysis, restricted to genes whose computed median expression among the 29 patients was at least 10 reads per million in both protocols. After computing the gene specific difference in the expression from the total RNA and the poly(A) counts (i.e., “differential expression’’, which measures the technical difference), we added this difference to the expression computed from the poly(A) counts. As can be seen from Fig. 1G and 1H, after such correction, the expression of coding and annotated non-coding RNAs perfectly clusters according to the patient the sample was derived from, unlike repetitive elements which are still not captured by poly(A) sequencing (Fig. 1I).

### Expression of repeats exhibits higher noise than that of the conventional genes

We tested the effect of technical noise on whether the bias of the computed gene expression is protocol specific and gene independent. As we have mentioned, in the absence of technical noise one would expect the ratio of expression between the poly(A) and total RNA protocol to be a sample independent constant for each gene. We performed the chi-squared test for the variance of the ratio of the computed expressions for each sample. We required that the variance of this ratio across the samples did not exceed the biologically significant expression difference. This would imply that the biologically significant change is higher than the technical noise; and meaningful differential expression is detectable using the smallest number of biological replicates. As a result, 61% of the coding genes, 37% of annotated non-coding RNA and only 8% of Repbase elements passed the test at the FDR cutoff of 0.05. Genes and repeats that did not pass the test would require a larger sample size to detect the biologically significant effects.

We further measured the rank correlation between expressions of each gene and repeat element detectable using both protocols (Fig. 2A). At the FDR value of 0.05, 99% of the coding genes, 95% of the annotated non-coding RNA and 56% of repeats passed the test. As can be seen from Fig. 2A, expression of repetitive elements exhibit a positive correlation; however, the value of this correlation is smaller than for the coding and annotated non-coding RNA. The reason for this may be technical noise caused by the effect of DNA contamination being stronger for repeats since their loci in the genome are typically much longer than those of the coding/annotated non-coding RNA. Alternatively, some repeats with a poly(A) tail may have additional copies in the genome lacking poly(A) tail, making the transcripts of those extra copies undetectable by the poly(A) protocol. On the other had, all mRNA have a poly(A) tail.

**Figure 2.**
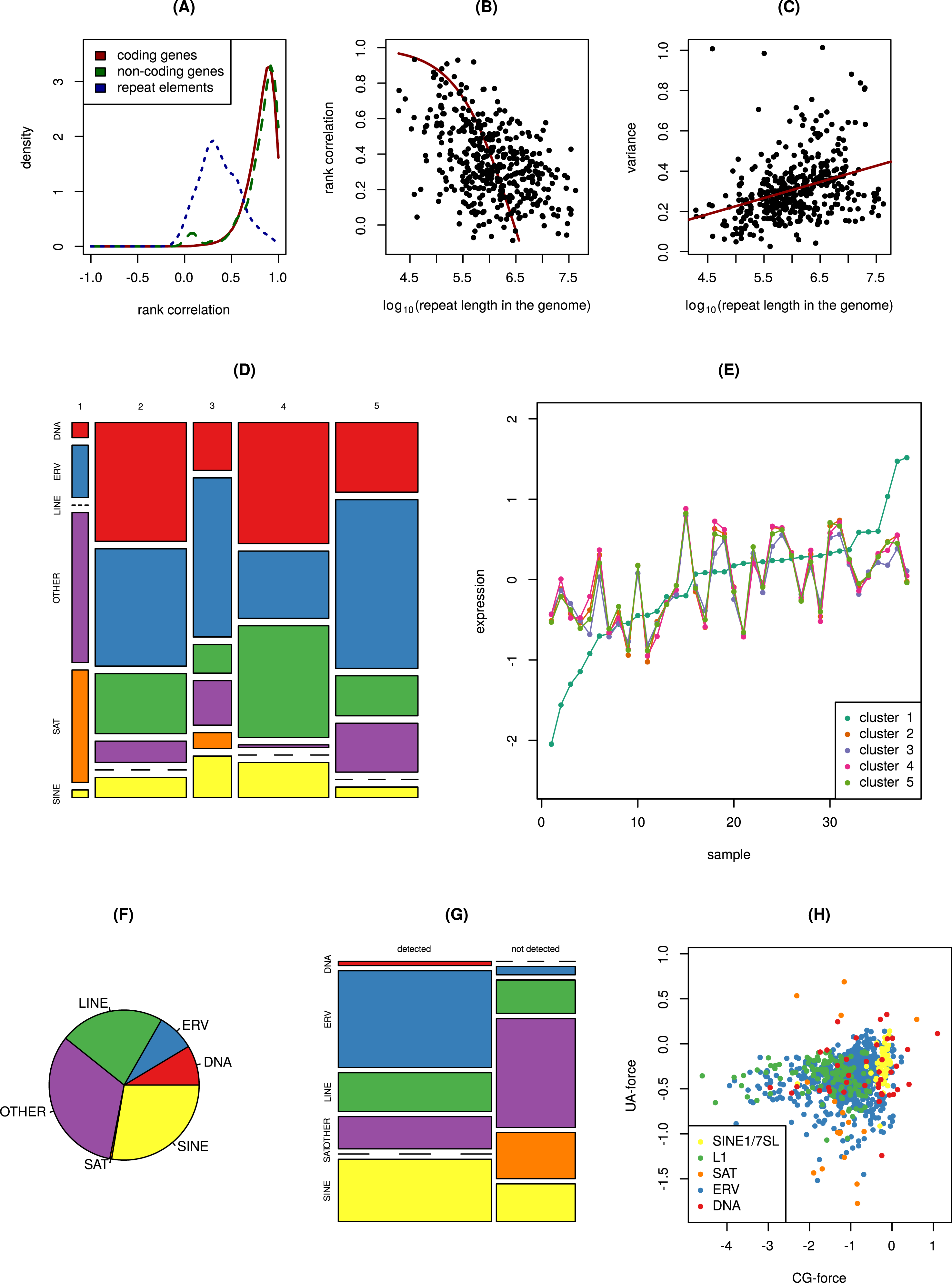
Rank correlation between the expression according to the total RNA and the poly(A) data was computed for each gene and repeat element detectable using both protocols. Distribution of the rank correlation for the coding and non-coding genes as well as repeat elements is shown. Rank correlation of repeat expression is typically smaller than that of the coding or non-coding genes since repeats experience a higher technical noise (t-test, p=3e-168). Small peak for the noncoding genes near zero comes from rRNA. See also Table S3. Regression for the rank correlation between the repeat expression according to the total RNA and the poly(A) data vs. length of the integration sites of the repeat element in the genome. Repeats with a higher length have smaller correlation. Linear regression for the variance of the computed expression difference for each repeat element vs. length of the integration sites of the repeat element in the genome. Repeats with a higher length have higher variance. See also Table S2. Cluster assignment vs repeat type. See also Table S4, Fig. S1 (B, C). Consensus (median) expression within the five repeat clusters. Proportion of different repeat types within repeat reads. Here we have not included the counts for rRNA, pseudogenes and snRNA. See also Fig. S1 (A). Detectability of repeat elements of different types in the poly(A) RNA-seq. Note that the satellites (SAT) are not detectable, and DNA transposons (DNA) are detectable. Most of the ERV/LINE1/SINE are detectable. See also Fig. S1 (C). CG- and UA- compositional bias computed for the consensus sequence for repeats of different types. Satellites (SAT) exhibit the highest diversity.

We investigated the connection between correlation and cumulative length of a repeat sequence within the HG38 genome. These values are negatively correlated (rank correlation rho=-0.42, p=8e-19). We performed regression between these variables (see Fig. 2B, p-value < 2e-16), which predicted perfect correlation between expression values computed using the two protocols is achieved for a cumulated sequence length of 13 kilobases. Regression between the variance of expression difference between the two protocols and cumulative length of repeat sequences (Fig. 2C, p-value=1.29e-12), further supports the observation that repeats with a higher length of integration sites within the genome exhibit greater noise. This is consistent with genomic DNA contamination as the cause of variation.

### Repetitive elements form distinct co-expression clusters

We performed consensus clustering of repetitive elements using the 39 total RNA tumor samples in TCGA (29 of which have paired poly(A) select samples). Five clusters of repetitive element coexpression were detected, indicating many repetitive elements aberrantly expressed in tumors are not independently expressed of one another, but are co-expressed (Fig. 2D and 2E). Such clustering further indicates different clusters of repeat expression may confer different phenotypic traits. One cluster is an outlier in terms of its expression and contains most of the satellite repeats (Fig. 2D, 2E). This cluster exhibits the highest diversity of expression across tumors, implying that satellite repeats are most likely to have individualized patterns of expression. Similar behavior was observed in earlier datasets of mostly pancreatic cancer samples performed on a different sequencing platform (Ting et al. 2011). The other four clusters involve respectively LINE, SINE, ERV, and DNA and other repeats labeled as “Other” (e.g, CR1, hAT, simple repeats) (Fig. 2F). Unlike the cluster containing most SAT repeats, these clusters have similar consensus expression. The reason for their existence as separate clusters is that we use centroid based clustering, which is prone to splitting clusters with many similar (overrepresented) elements, which is particularly the case for LINE and SINE elements.

We compared detectability of each repetitive element class by the poly(A) protocol (Fig. 2G). Strikingly, contrary to ERV, LINE and SINE, satellite repeats appear almost universally undetectable by this poly(A) protocol, despite studies reporting that a fraction of these transcripts are actively polyadenylated (Criscione et al. 2014). This highlights the importance of total RNA-seq to further study the role of satellite repeats in cancer, particularly their role in initiating the innate immune response in the tumor microenvironment (Tanne et al. 2015). Of note, the raw read number of SAT repeats comprises only a small fraction of all of repeat reads, typically less than 1%.

In an earlier study, immunostimulatory properties of aberrantly expressed repeats was ssociated with unusual usage of dinucleotide motifs compared to the rest of the human genome (Tanne et al. 2015). We therefore quantified aberrant motif usage by the forces on CpG and UpA, dinucleotides, which measure their deviation from maximum entropy dinucleotide usage values. We computed these effective forces for all LINE, SINE and SAT elements (Fig. 2H), see (Tanne et al. 2015) and (Chatenay et al. 2017) for an overview of the methods. Interestingly, in agreement with (Tanne et al 2015), satellite elements are the most diverse in terms of the CpG and UpA compositional bias, and consequently we proposed they are more likely to engage immune receptors such as pattern recognition receptors (PRRs) (Vabret et al. 2017).

### ERV class expression is associated with positive anti-PD-Ll immunotherapy response

Pre-existing tumor T cell inflammation can be a strong predictor of response to cancer immunotherapy such as anti-PD-L1/PD-1 or anti-CTLA-4 antibodies (Chen et al. 2017). Several studies have recently highlighted links between tumors ERV expression, the expression of “viral defense genes”, and anti-tumor responses (Chiappinelli et al. 2015, Roulois et al. 2015, Badal et al. 2017). It was hypothesized that chemically-induced epigenetic dysregulation in tumors leads to expression of ERVs, which in turn stimulate innate immune PRRs and create an anti-tumoral innate immune response. In one of these studies (Chiappinelli et al. 2015), this response was associated with clinical benefit in patients treated with anti-CTLA-4 therapy. We examined one of the few available tumor immunotherapy RNA-seq datasets from patients treated with PD-L1 blockade (Snyder et al. 2017). In this cohort of patients with urothelial cancer, we tested the hypothesis that ERV expression is also associated with clinical benefit from therapy. We performed this analysis for the first time in an anti-PDL1 treated tumor, as opposed to the previous anti-CTLA4 studies.

We performed hierarchical clustering using expression of ERV repeats using the repeatmasker/Repbase annotation, which revealed two distinct clusters of high and low ERV expression levels (Fig 3A). In this case, association between ERV repeats expression and patient response to PD-L1 immunotherapy was significant (p=0.024, Fisher’s exact test). Consequently, patient survival analysis showed that high expression of ERV repeats correlates with overall survival (Fig. 3D, p=0.012) and progression free survival (Fig. 3E, p=0.025). Interestingly, expression of ERV repeats was a better predictor of response to immunotherapy than the viral defense signature in this cohort, which did not similarly segregate patients (Fig. 3C). Additionally, as we show that Repeatmasker/Repbase annotation for ERV repeats yields a higher read number than that for ERV genes annotated in Ensembl, we suggest that clinical studies would reveal more accurate associations by interrogating global repeat expression rather than specific ERV genes as those annotated in Ensembl, or associated immune classes. It is worth mentioning that the read counts of the ERV genes annotated in Ensembl were below the standard 10 reads per million threshold in RNA-Seq, ERV3 and ERV3K having the highest read number. Expression of these two genes is correlated with the mean ERV expression (Fig. 3B). The implication is that, due to the abundant transcription of repetitive elements, they are a more robust predictor of response to immunotherapy than the expression of associated immune genes, which likely require a larger sample size to resolve cohorts.

**Figure 3.**
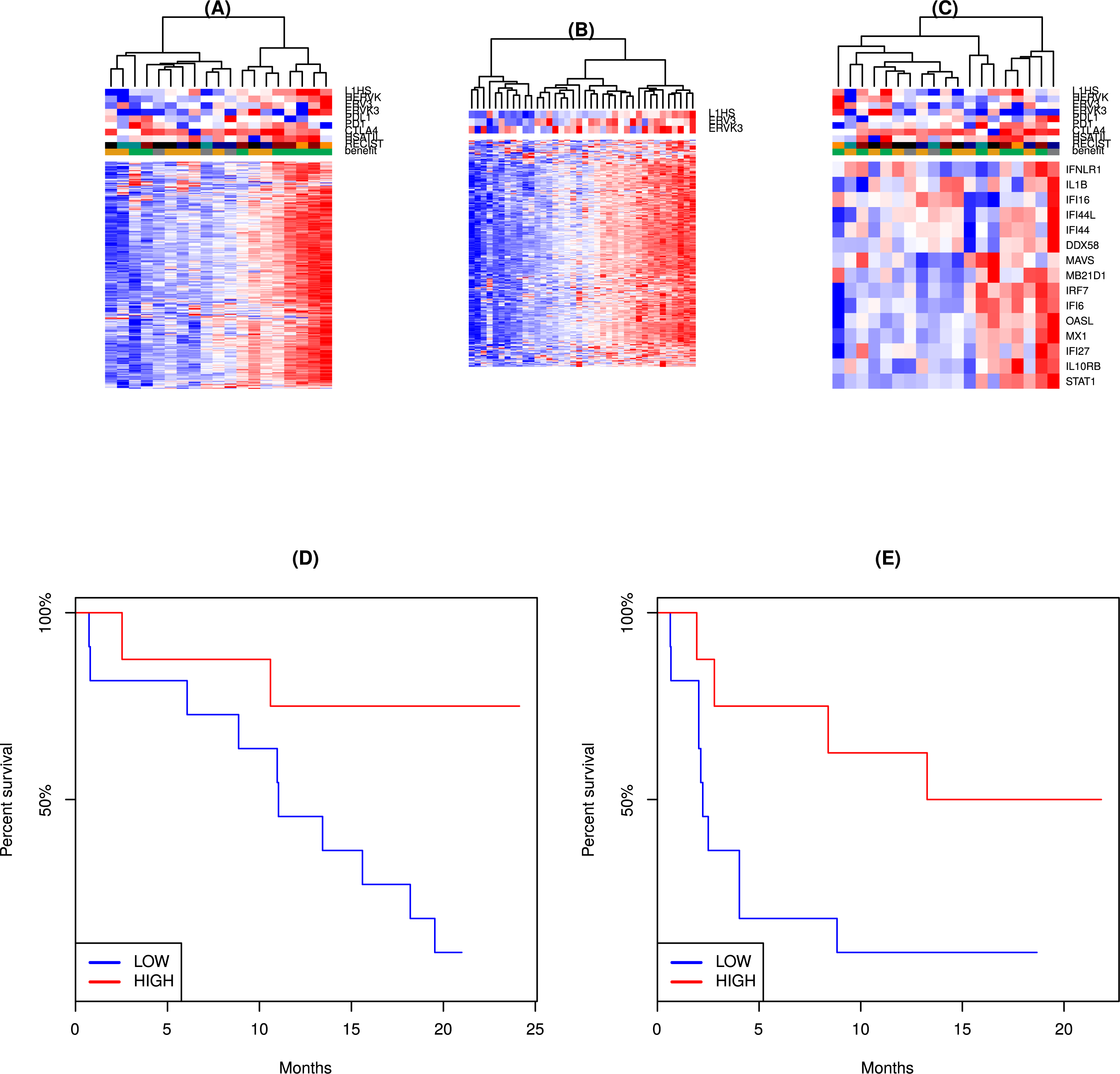
Heatmap for the ERV repeat expression in urothelial cancer dataset from Snyder et al. Annotation at the top: L1HS, HERVK, HSATII - expression of the corresponding repeat elements. ERV3, ERVK3, PD-L1, PD1, CTLA-4 - expression of the corresponding Ensembl genes. The read counts for ERV3 and ERVK3 are the highest among ERV genes annotated in Ensembl; nevertheless, they are still below the conventional low bound in RNA-Seq (10 reads per million) in all samples. RECIST: black - missing data, blue - PD (progressive disease), cyan - SD (stable disease), orange - PR (partial response), red - CR (complete response). Benefit: green - clinical benefit, orange - no clinical benefit, gray - long survival despite the absence of the clinical benefit. Heatmap for the ERV repeat expression in the TCGA total RNA dataset. Annotation at the top: L1HS - expression of the corresponding repeat element. ERV3, ERVK3 - expression of the corresponding Ensembl genes. Pearson correlation between the mean expression of ERV elements and expression of ERV3 gene is 0.46 (p=0.0040, t-test, two-tailed). Pearson correlation between the mean expression of ERV elements and expression of ERVK3 gene is 0.40 (p= 0.013, t-test, two-tailed). Heatmap for the interferon stimulated (viral defense) gene expression in urothelial cancer dataset from Snyder et al. Color annotation at the top is the same as that in Fig. 3A. Kaplan-Meier plot for the overall survival between the patients from the ERV repeat high and ERV repeat low clusters. Association is significant (p=0.012, log rank test). Kaplan-Meier plot for the progression free survival between the patients from the ERV repeat high and ERV repeat low clusters. Association is significant (p=0.025, log rank test).

### Global repeat depression is associated with an immunosuppressive phenotype

We next studied the relation between expression of repetitive elements and tumor progression in human cancers not treated with immunotherapy. As few total tumor RNA-seq data are publicly available, we examined the expression of LINE and ERV elements, which can be detected using poly(A) capture, increasing our sample size. We focused on LINE and ERV expression in colon and rectal adenocarcinoma cancers available in TCGA, given the well-established genetics of colon cancer progression, the established co-expression of LINE1 and HERV-K (Desai et al. 2017), and the known presence of satellite repetitive element genome expansions (Bersani et al. 2015). We examined 364 paired-end RNA-seq samples prepared with the poly(A) protocol. We first sorted samples by their expression level of LINE1 elements most recently integrated into the genome (L1HS) and performed differential expression analysis between the third and the first tercile. Survival analysis (Kaplan-Meier curve) using the TCGA data shows that patients from the lowest L1HS expression tercile have longer survival, compared to the patients from the highest L1HS expression tercile (p=0.0297) (Fig. 4A). Expression of LINE1 may signal an advanced loss of epigenetic control that correlates with tumor progression and influences patient prognosis.

**Figure 4.**
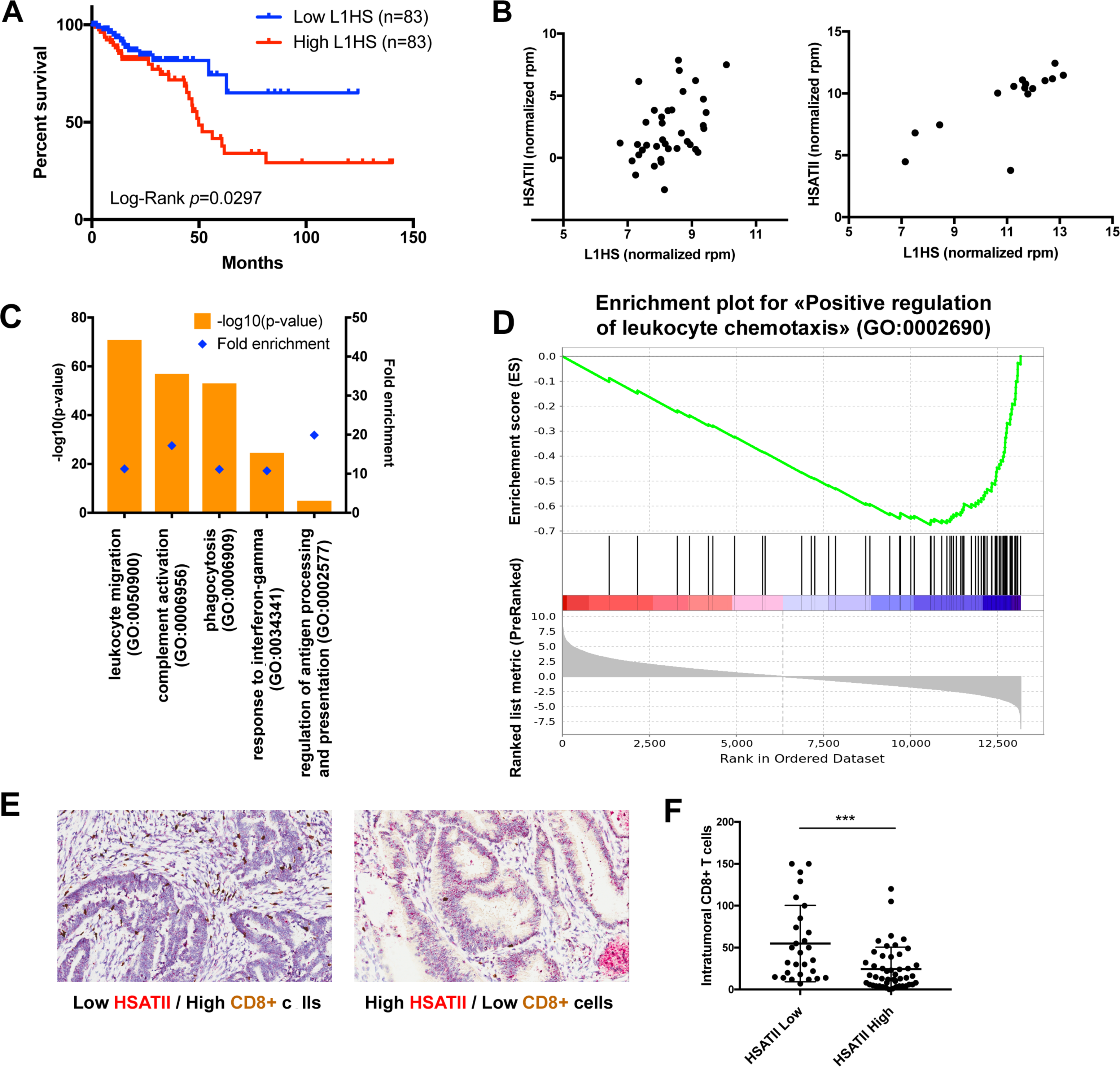
Kaplan-Meier plot depicting the survival over time for patients with high (red - top tercile) and low (blue - bottom tercile) L1HS expression. Dataset comes from colon and rectal adenocarcinoma cancers available in TCGA and classified as microsatellite-stable. See also Table S5, S6. Correlation of HSATII and L1HS expression in tumors prepared with total RNA protocol available in TCGA (n=38, left) and in pancreatic tumors sequenced by single-molecule sequencing (n=16, right). GO terms enriched in genes downregulated in the third compared to the first tercile of samples sorted by L1HS expression in TCGA MSS colorectal tumors. See also Table S5 and S6. GSEA enrichment plot for genes of the “Positive regulation of leukocyte chemotaxis” GO set. Genes were ranked by the t-statistic produced by comparison of their expression in the third and the first tercile of samples according to L1HS expression in TCGA MSS colorectal tumors. P-value p<1e-4. Representative images of colon tumor stained for CD8 protein expression (immunohistochemistry - brown) and HSATII RNA (in situ hybridization - red). Left picture: low HSATII expression correlates with high CD8+ T cells infiltration. Right picture: high HSATII expression correlates with low CD8+ T cells infiltration. Associated quantification of colon cancer intratumoral CD8+ T cell per field of view (400×200μm). Tumor samples were classified as HSATII high or low expression following in situ hybridization staining. p-value=0.0004 (unpaired t-test). See also Table S7.

To study in detail the relationship between repeat expression and cancer progression, we further analyzed the difference in gene expression in tumors expressing high or low levels of human LINE1. Gene ontology (GO) enrichment analysis uncovered significant enrichment of specific GO terms when analyzing the subset of genes downregulated in high versus low LINE1 expression samples. Interestingly, all the terms were related to immune response, suggesting they are the main pathways associated with LINE1 expression. Moreover, the samples that show upregulation of LINE1 expression demonstrated no significantly enriched GO term. The most significant GO terms enriched for the downregulated genes include *leukocyte migration, complement activation, phagocytosis, response to interferon-gamma* and *regulation of antigen processing and presentation* (Fig. 4C). We also performed GSEA analysis on one of the enriched GO terms, *positive regulation of leukocyte chemotaxis* (Fig. 4D). The implication is that either there is a correlation between the lack of epigenetic control associated with LINE1 expression and immune suppression, or, to the extent to which LINE1 elements engage immune pathways, they are activating pathways associated with negative regulation (See also Figure S1D).

As similar gene expression analysis could not be performed with satellite repeats due to the low number of total RNA sequence available, we measured the relationship between LINE1 and specific satellite RNAs. Previous work using single molecule RNA-seq had shown a strong association of LINE1 repeats with pericentromeric satellites in both mouse and human cancers (Ting et al. 2011). We confirmed LINE1 expression correlates with expression of the human pericentromeric satellite HSATII in the TCGA tumor samples prepared with total RNA protocol and in pancreatic tumors sequenced by single-molecule sequencing, obtained from (Ting et al. 2011) (Fig. 4B). Given the ability of single molecule RNA-seq to better quantify HSATII, we performed a targeted analysis of the 16 such pancreatic cancer samples (Ting et al. 2011) to determine if there was a consistent relationship between HSATII and the tumor immune microenvironment. We binned samples into terciles according to HSATII expression and performing differential expression analysis between the third and the first tercile. In particular, genes downregulated in HSATII high samples were also enriched in the *lymphocyte migration* GO term. Additionally, we performed a GO-independent analysis of immune gene enrichment following the immune signatures defined in (Rooney et al. 2015). Interestingly, the two genes labeled as responsible for the cytolytic activity (GZMA and PRF1) associated with cytotoxic T (CD8+) activation are highly downregulated in high HSATII expressing samples (8-fold change).

To validate the relevance of these GO terms, we performed combined RNA *in situ* hybridization for HSATII and immunohistochemistry for cytotoxic T cells (CD8+) in a cohort of 75 colon tumor samples (Fig. 4E, 4F). We scored tumors based on high or low levels of HSATII by comparing relative levels of HSATII staining in tumor cells compared to normal adjacent cells. We then quantified the density of CD8+ T-cells observed in the tumor microenvironment finding significantly lower CD8+ T-cells in HSATII high tumors. This is consistent with our computational analysis of RNA-seq data demonstrating a downregulation of immune related GO terms in repeat (LINE1 or HSATII) cancers. In previous studies, HSATII RNA demonstrated direct immunostimulatory properties of dendritic cells through a Myd88-dependent pro-inflammatory cytokine response (Tanne et al. 2015). Interestingly, the anti-correlation we observed between HSATII expression and tumor lymphocyte infiltration could suggest a complex mechanism where HSATII mediated signaling would be causally linked to a protumoral anti-inflammatory response that is associated with CD8+ T cell exclusion. This would be consistent with previous work demonstrating that specific stimulation of innate immune receptors on cancer cells can be protumorigenic, such as in pancreatic cancer (Zambirinis et al. 2014) where HSATII is known to be highly abundant (Ting et al. 2011). Therefore, it can be hypothesized that HSATII-mediated signaling, largely undetected by poly(A) RNA-seq protocols, could be partially responsible for inducing an immunosuppressive microenvironment preventing lymphocyte infiltration. In both cases, these observations call for a broader survey of HSATII, LINE1 and other repeat RNAs in cancer using total RNA-seq protocols.

## Discussion

Broader use of total RNA sequencing protocols and single molecule sequencing platforms would allow researchers to investigate the expression of repetitive elements and their use as biomarkers or immune stimulators in cancer. Available data reveal that the conventional poly(A) capture based RNA sequencing allows one to detect expression of only a limited number of repetitive elements, despite their recently established role in prognosis and response to epigenetic and immunotherapy. Only a subset of LINE, SINE, and ERV related elements can be captured with the poly(A) protocol, along with some DNA repeats. Conversely, satellite repeats (in particular, HSATII, a known cancer biomarker and immunostimulatory molecule) are only detected using the total RNA protocol.

The relationship between the expression of most endogenous retroviruses, LINE and SINE elements indicates that they use a similar biological mechanism for transcription, which may be decoupled from satellite repeat transcription. However, one will need larger sample sizes in order to adequately quantify and study the variability of the expression of repetitive elements as compared to the coding genes and annotated non-coding RNA. The utility of total RNA is evident for non-coding RNA. Additionally, a stranded preparation protocol is required for testing the established hypothesis that RNA, transcribed from both sense and antisense strands form long double-strands that may activate viral patterns recognition receptors (PRRs).

In previous studies ERV expression has been linked to an anti-tumoral response in very early stage primary melanoma (Badal et al. 2017), and a positive response to epigenetic and anti-CTLA-4 immunotherapies (Chiappinelli et al. 2015, Roulois et al. 2015). We show that ERV expression is associated with positive response in a set of patients treated with anti-PD-L1 therapy, extending previous findings in melanoma patients treated with anti-CTLA-4. Moreover, while ERV expression segregated patients, the viral defense signature associated with response in previous work did not, suggesting that abundant transcription of repetitive elements may offer a more robust biomarker.

Satellite repeats display heterogeneous expression and anomalous motif usage relative to other repeat classes, however only LINE1 and ERV classes can be probed adequately using poly(A) protocols. Based upon this one may expect that repeats are generally better for prognosis. However, here we find their expression is linked to poorer prognosis in typically late stage pancreatic and colon cancers. In colon cancer, LINE1 element expression correlates with lack of cytotoxic T-Cell infiltration using poly(A) select samples. This presents a curious paradox. A possible resolution is that LINE1 expression is correlated with many other repeats as can be seen from total RNA-seq; in particular the satellite repeat HSATII, since LINE1 is essentially expressed whenever HSATII is expressed, though not vice versa. The implication is that in the studies where LINE and ERV expression was anti-tumoral, molecules such as HSATII were not additionally coexpressed. Indeed in the study of Badal, et al., which used the total RNA protocol and found a positive association between repeat expression and prognosis, little HSATII expression was observed in these early stage tumors. Consistent with this finding, lower mouse satellite repeat expression was also found in early versus advanced pancreatic tumors in a genetically engineered mouse model using RNA-ISH (Ting et al. 2011). It may be the case that in late stage tumors, where abundant repetitive element expression is associated with failure of tumor suppressors, the large scale transcription of many “non-self” repetitive elements has been coopted by the tumor’s evolution to maintain an advantageous inflammatory state. The distinct sequence motifs in satellite RNAs including HSATII that appear “non-self” leading to differential innate immune response is consistent with this theory (Tanne et al PNAS 2015). Altogether, this indicates that response to repeat RNAs are heterogeneous, which leads to relative changes in the balance of inflammatory immune response that are pro- or anti- tumoral.

To support this hypothesis, we demonstrate in our study the association of HSATII RNA expression with lack of CD8 T-cell infiltrates and suggest that its expression may induce a response immunosuppressive to cytotoxic T-cell infiltration, and therefore is a confounding factor in studies which only monitor LINE and ERV expression. Since HSATII is not detected by poly(A) sequencing protocol, we conclude that causal molecules with a critical role engaging the innate response in the tumor microenvironment, may be hidden from view using current sequencing protocols. We argue that HSATII functions as a novel epigenetic checkpoint in the tumor microenvironment, underappreciated due to its lack of sequence visibility, and initiating cytokine signaling in an pro-tumoral manner. Indeed TLR mediated signaling, which HSATII has been shown to engage (Tanne et al. 2015), has recently been implicated in pro-tumoral inflammation in pancreatic cancer (Ochi et al. 2012), where tumors have learned to express single-stranded RNA sensors, such as TLR7. Curiously, HSATII expression is abundant in this same tumor type (Ting et al. 2011). As a result we demonstrate the need for total RNA protocols and associated bioinformatics tools to discover currently hidden, yet likely critical, signaling RNAs in the microenvironment.

## Experimental Procedures

We have selected all samples from TCGA which had both total RNA and poly(A) enriched RNA-seq data derived from different aliquots of the same physical sample. In total there are 29 such samples. After initial quality filtering we aligned the reads to the human genome and to the Repbase database of repetitive elements (Bao et al. 2015).The number of reads mapping to the annotated genomic features was quantified and expression was computed.

Additional details of the analyses are given in the Supplemental Experimental Procedures.

## Author contributions

AS, KSA, BDG and DTT conceived the study and design. AS, NV, KSA, NB, RM, SC, BDG and DTT developed methodology. AS and KSA acquired data. AS, NV, KSA, RM, SC, A Snyder, SF, DB, JR, BDG and DTT analyzed and interpreted data. AS, NV, KSA, NB, RM, SC, BDG and DTT wrote, reviewed, and/or revised the manuscript. BDG and DTT supervised the study.

## Acknowledgements

This work was supported by the NIH P01CA087497-1 (BDG), Stand Up to Cancer (DTT, BDG), the Lustgarten Foundation (DTT, BDG), the National Science Foundation 1545935 (DTT, BDG), the Burroughs Wellcome Fund (DTT), and Affymetrix, Inc. (KSA, DTT). AS and BDG would like to thank Remi Monasson and Simona Cocco for many helpful comments and suggestions.

